# The gap junction blocker mefloquine impairs sleep-dependent declarative memory consolidation in humans

**DOI:** 10.1101/868901

**Authors:** Gordon B. Feld, Hong-Viet Ngo, Ernesto Durán, Sandra Gebhardt, Lisa Kleist, Kerstin Brugger, Andreas Fritsche, Jan Born, Manfred Hallschmid

## Abstract

During sleep, the time-compressed replay of engrams acquired during preceding wakefulness drives memory consolidation. We demonstrate in healthy humans that direct electrical coupling between neurons via gap junctions, i.e., electrical synapses, contributes to this beneficial effect of sleep. Twenty male participants learned a declarative word-pair task and a procedural finger sequence tapping task before receiving the antimalarial mefloquine that is known to block electrical synapses. Retrieval was tested after a retention interval of approximately 20.5 hours that included nocturnal sleep. As predicted, mefloquine given before sleep impaired the retention of declarative memory. In contrast, this effect was absent in control groups, which stayed awake or received mefloquine after sleep. Irrespective of sleep or administration time, mefloquine enhanced retention performance on the procedural memory control task. We conclude that sleep-dependent processes relying on electrical neuronal coupling enable hippocampus-dependent declarative memory consolidation, presumably via time-compressed hippocampal replay of memory traces within sharp-wave/ripple complexes. The recruitment of this understudied form of neuronal information transfer may be necessary to achieve fast-paced memory reprocessing during sleep. Considering that drugs targeting neurochemical synapses have recently fallen short of substantially advancing the treatment of memory impairments in Alzheimer’s disease, schizophrenia or during normal aging, unraveling the contribution of gap junctions to sleep-dependent declarative memory formation can be expected to open new therapeutic avenues.

**Significance statement:** Sleep supports the strengthening and transformation of memory content via the active replay of previously encoded engrams. Surprisingly, blocking neurochemical synaptic transmission does not impair this function of sleep. Here we demonstrate that the direct electrical coupling between neurons via electrical synapses (gap junctions) is essential for the sleep-dependent formation of declarative memory, i.e., memory for episodes and facts. These findings are in line with the assumption that electrical synapses enable time-compressed neuronal firing patterns that emerge during sleep and drive declarative memory consolidation. Electrical synapses have so far not been linked to higher-order brain functions in humans; their contribution to sleep-dependent memory processing may provide a novel target for sleep-related clinical interventions.

## Introduction

Sleep is an efficient and selective memory enhancer (Stickgold, 2005; Rasch and Born, 2013; Feld and Born, 2019), but its effectiveness is subject to the amount of information learned before sleep (Feld et al., 2016). Accordingly, when memory traces that were acquired during wakefulness are replayed during subsequent sleep this replay is compressed in time by a factor of 5-10 (Ji and Wilson, 2007), which greatly enhances the amount of information that can be processed. Replay stabilizes memory representations, thereby consolidating the initially labile traces into lasting memories (Rasch et al., 2007; Dupret et al., 2010; Yang et al., 2014), and occurs mainly during hippocampal sharp-wave/ripple events (Diba and Buzsaki, 2007). Consequently, suppressing these events electrically or optogenetically impairs memory retention (Girardeau et al., 2009; van de Ven et al., 2016). Preserved sequential firing during replay, as well as the electrophysiological properties of sharp-wave/ripples, suggests that neuronal plasticity emerging during sleep relies on Hebbian processes such as spike timing-dependent plasticity (Sadowski et al., 2016). However, blocking AMPA or NMDA receptor-mediated synaptic signaling, i.e., the foremost neurochemical processes generating such plasticity, does not compromise the contribution of sleep to declarative memory consolidation (Feld et al., 2013), leaving open the question which neuromolecular mechanisms coordinate sleep-induced plasticity.

Here we investigated whether the direct electrical coupling of neurons via gap junctions, i.e., electrical synapses, is more relevant for these processes. Gap junctions formed by connexins nesting on neighboring hippocampal neurons enable fast network oscillations via direct electrical coupling (MacVicar and Dudek, 1981; Draguhn et al., 1998; Schmitz et al., 2001) and contribute to the generation of sharp-wave/ripple complexes (Maier et al., 2003) that mark memory replay in the hippocampus (Diba and Buzsaki, 2007). In-vitro evidence also indicates that mefloquine blocks very fast hippocampal oscillatory firing (300 Hz) during recurrent epileptiform discharges (Behrens et al., 2011), but this effect may also be attributable to nonspecific drug effects. Since this existing evidence from animal models putatively links gap junctions to sleep’s facilitating effect on memory consolidation (Stickgold, 2005; Rasch and Born, 2013) and especially to memory-enhancing features of sleep electrophysiology (Staresina et al., 2015), we decided to directly test the hypothesis that electrical synapses play a major role in processes of memory formation in the offline brain.

To this end, we chose mefloquine to selectively block gap junctions containing the connexin 36 channel (Cruikshank et al., 2004) during a retention interval that included eight hours of nocturnal sleep. Before administration participants learned a declarative word-pair task and a procedural finger sequence task, which were both retrieved on the next day. We predicted that mefloquine would selectively disrupt sleep-dependent declarative memory consolidation that heavily recruits the hippocampus, while not affecting sleep-dependent procedural memory consolidation that relies on other structures (Zola-Morgan and Squire, 1990). We performed two additional control experiments to exclude similar effects of blocking gap junctions during wakefulness or at retrieval.

## Materials and Methods

### Participants

Our placebo-controlled counter-balanced within-subject study included a total of forty-four participants (Experiment 1: n = 20, Experiment 2: n = 12, Experiment 3: n = 12). They were healthy, non-smoking, native German-speaking men (18-30 years) who held degrees qualifying them for secondary education and were not taking prescribed medication. None of them reported any past or present chronic physical or psychological illness, and routine examinations prior to inclusion confirmed the absence of any acute mental or physical disease. Participants reported a normal sleep-wake pattern during the week preceding each session, did not work night-shifts and had not traveled across more than six time zones in the twelve weeks before participation. Participants were instructed to get up at 7:00 am on the days of the experiments and not to ingest alcohol or (after 1:00 pm) caffeinated beverages. In Experiments 1 and 3, an adaptation night was conducted at least 48 hours before the actual experiment under identical conditions (i.e., including electrodes for polysomnographic recording). The experiments were approved by the local ethics committee (Ethics Committee of the Medical Faculty at the University of Tübingen). Written informed consent was obtained from all participants before participation and they were compensated financially.

### Experimental design and procedures

All three experiments followed a double-blind, placebo-controlled, within-subject, balanced crossover design (see Figure 1A for an overview). Each experiment consisted of two experimental sessions that were conducted at least four weeks apart and used an identical set-up including parallel versions of the learning tasks. In counter-balanced order, participants received 250 mg mefloquine (Lariam ®, plasma peak: 10 hours, half-life: 10 days, see Gbotosho et al., 2012) in one and placebo in the other session. Identical capsules containing mefloquine or placebo and manufactured according to GMP were prepared by the pharmacy at the University Medical Centre of the University of Mainz, Germany.

**Figure 1.**
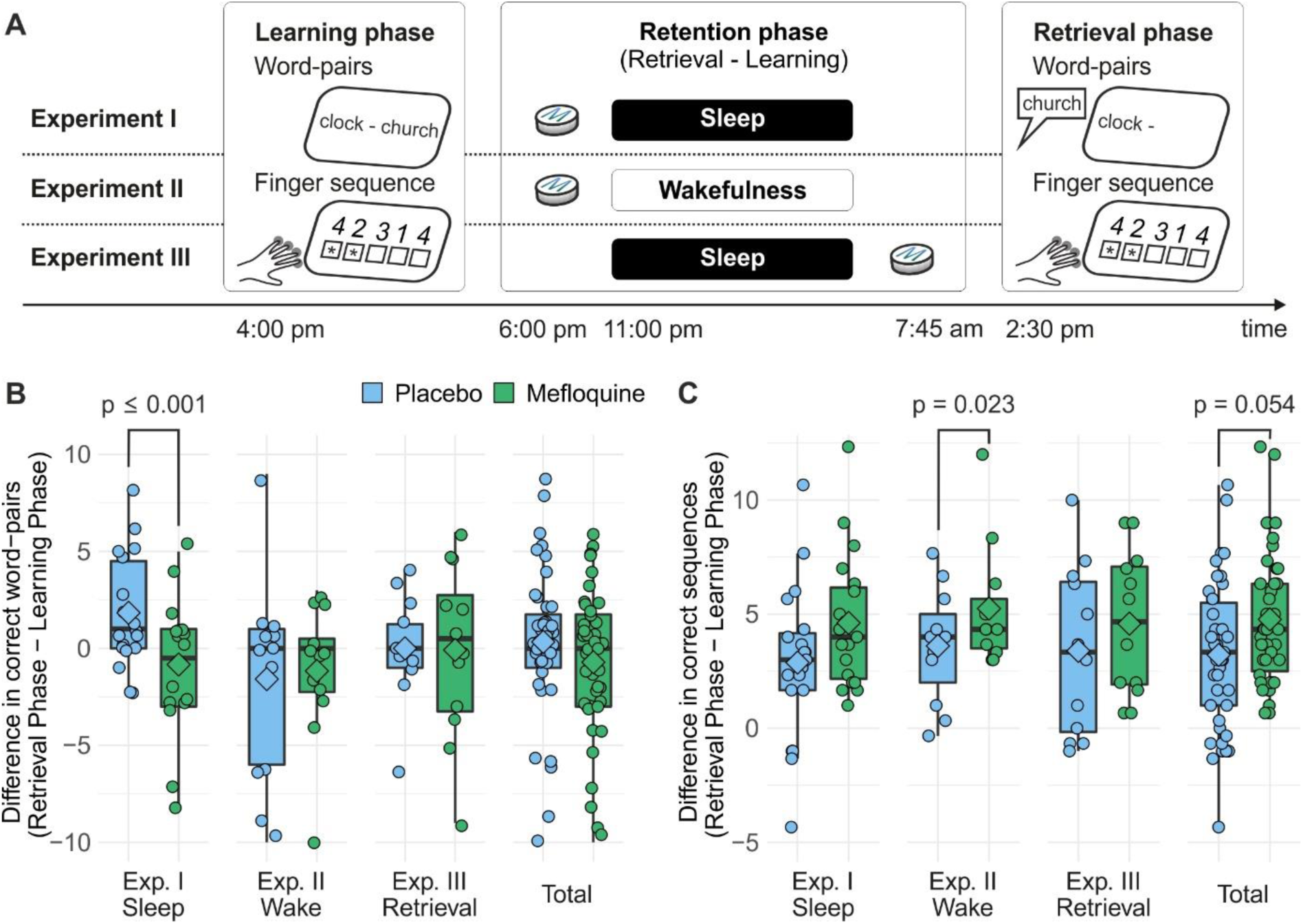
Experimental design and results of memory retention. (A) In three experiments we administered the gap junction blocker mefloquine to healthy humans in order to suppress direct electrical coupling between neurons and assess effects on sleep-dependent memory consolidation. Participants learned a declarative word-pair task and a procedural finger sequence tapping task in the afternoon. Retrieval was tested in the subsequent early afternoon. To test declarative memory formation, forty word-pairs were first presented to and then immediately recalled by the participants in response to the first word. Participants received instant feedback and recall was repeated until they achieved 60% correct answers. This procedure was repeated during retrieval without feedback; the absolute difference between immediate recall during the learning and recall during the retrieval phase represented declarative memory retention. For the assessment of procedural memory, after the word-pairs, a five-digit sequence was displayed, and participants were asked to tap it with the fingers of their non-dominant hand. They did so during twelve blocks of 30-second tapping and 30-second break intervals. During retrieval they performed an additional three blocks. The difference between the average amounts of correctly tapped sequences during retrieval and the last three blocks of learning represented procedural memory retention. In Experiment 1, participants received 250 mg of mefloquine (or placebo) at 6:00 pm to induce effective gap junction blockade during sleep. In Experiment 2, participants received the drug at the same time, but were not allowed to sleep until after retrieval. In Experiment 3, participants were allowed to sleep, but received the drug after sleep to block gap junctions at retrieval only. (B) Box and whisker plot (thick line: median, hinge: 25% and 75% quartiles, whiskers: smallest/largest values no further than 1.5 * inter quartile range from the hinge), mean (diamond shape) and individual data points (circles) of retention performance for word-pairs for the mefloquine condition (green) and the placebo condition (blue) in the individual experiments and across all three experiments (Total). As hypothesized, mefloquine significantly reduced the amount of word-pairs that participants retained only when it was active during sleep (Experiment 1: F_(1,16)_ = 29.75, p ≤ 0.001, η_p_^2^ = 0.65), indicating that gap junctions contribute to the sleep-dependent consolidation of declarative memory. (C) Box and whisker plot (thick line: median, hinge: 25% and 75% quartiles, whiskers: smallest/largest values no further than 1.5 * inter quartile range from the hinge), mean (diamond shape) and individual data points (circles) of retention performance in the finger-sequence task for the mefloquine condition (green) and the placebo condition (blue) in the individual experiments and across all three experiments (Total). Unexpectedly, mefloquine enhanced retention of the finger sequence irrespective of experimental group (F_(1,38)_ = 3.97, p = 0.054, η_p_^2^ = 0.10).

In general, participants arrived at the lab at 4:00 pm to start the learning phase (see *Memory tasks* for details). They always first learned word-pairs and then finger sequence tapping with ten minute-breaks of playing a computer game (available at Snood.com) in-between. After learning, vigilance, mood and sleepiness were assessed (see *Control measures*) and the participant received a snack (a traditional Swabian pretzel with butter) to enhance tolerability of mefloquine intake and watched animal documentaries (BBC’s ‘Planet Earth’). In Experiments 1 and 2, the drug was administered at 6:00 pm, thus ensuring sufficient blood concentrations at the beginning of the sleep or sleep-deprivation period (Gbotosho et al., 2012) without scheduling the learning phase too far from sleep onset time in the sleep group (Gais et al., 2006). Participants afterwards continued watching documentaries and, at 7:00 pm, received a standardized dinner (two slices of bread, butter, cheese and/or ham, tomato and decaffeinated tea).

Electrodes for polysomnography were applied at 9:00 pm in Experiments 1 and 3. At 10:15 pm blood was sampled for the assessment of mefloquine and the participant filled in questionnaires on mood and sleepiness. In these experiments, the electrodes were connected to the amplifier and lights were turned off at 11:00 pm. In Experiment 2, participants remained awake throughout the night and watched animal documentaries (see above). Participants of Experiments 1 and 3 were woken up between 6:45 and 7:15 am. Participants of all experiments filled in questionnaires on mood and sleepiness and blood was sampled at 7:30 am. Participants of Experiment 3 received the drug at 7:45 am (to induce high concentrations at retrieval of the memory contents). All participants left the lab in the morning and spent the time until retrieval as they pleased. They were however instructed not to sleep, to study or to consume caffeine or alcohol. Participants of Experiment 2 were continuously accompanied by an investigator to ensure wakefulness. They all returned to the lab at 2:30 pm and the memory tasks were retrieved in the same order as during learning (word-pairs and then finger sequence). Afterwards, participants performed memory control tasks (novel finger sequence tapping and number encoding, see *Memory tasks* and *Control measures*, respectively). Subsequently, general retrieval function, vigilance, mood and sleepiness were assessed, and the participant was asked if he assumed to have received mefloquine or placebo. Finally, a last blood sample was collected. We did not find indicators in any of the experiments that participants were able to identify the condition they were in (mefloquine or placebo; all p ≤ 0.34, exact McNemar’s test).

### Memory tasks

#### Word-pairs

The word-pair task to assess declarative memory performance robustly reflects sleep-dependent memory consolidation (Ekstrand et al., 1977; Plihal and Born, 1997). Participants started learning by seeing each of the 40 word-pairs for four seconds with an inter-stimulus-interval (ISI) of one second. Immediately afterwards, recall was tested in a cued recall procedure by showing the first word of each pair and asking the participant to say the matching second word. Irrespective of his answer, the participant afterwards received feedback on the correct word for two seconds. This cued recall procedure was repeated until the participant reached the criterion of 60% correct responses. During retrieval, the cued recall procedure was repeated without feedback. Performance at retrieval minus performance during learning represents retention.

#### Finger sequence tapping

The finger sequence tapping task measures procedural memory performance and shows robust sleep-dependent gains (Walker et al., 2003). During learning, participants tapped a five-digit finger sequence (e.g., 4-1-3-2-4) as fast and as accurately as possible during twelve 30-second blocks that were interrupted by 30-second breaks. To keep working memory demands at a minimum, the sequence was displayed on the screen at all times and an asterisk displayed under the current number was used to signal a button press. Responses were scored for speed (number of correctly completed sequences) and errors and this information was presented to the participant after every 30 second block. The final three blocks of the learning phase were averaged to indicate learning performance. During the retrieval phase, participants performed on an additional three blocks of the task and these were averaged to indicate retrieval performance. Performance speed was assessed as the total amount of correctly tapped sequences and performance accuracy as the error rate per block. Performance at retrieval minus performance during learning represents retention. After retrieval, participants performed on an additional three blocks of a new sequence to assess if potential performance changes at retrieval were due to memory-independent effects like unspecific changes in reaction speed.

#### Sleep scoring and frequency analyses

Standard polysomnography in Experiments 1 and 3 was performed using Ag-AgCl cup electrodes connected to a Brain Amp DC 32-channel EEG-amplifier (Brain Products GmbH, Gilching, Germany) for the EEG channels and a Brain Amp ExG 8-channel bipolar amplifier (Brain Products GmbH, Gilching, Germany) for the bipolar EOG and EMG channels that shared a ground electrode connected to the forehead. EEG was recorded from F3, Fz, F4, C3, Cz, C4, P3, Pz, P4 according to the international 10-20 system and referenced to coupled electrodes attached to the mastoids. Additionally, horizontal and vertical eye movements and muscle tone (from the chin) were recorded. Heart rate was recorded via ECG electrodes to monitor for adverse side effects. Data were low-pass- (80 Hz) and high-pass-filtered (0.16 Hz) before digitising at 250 Hz. For sleep scoring, they were down-sampled to 200 Hz and additional offline low-pass (35 Hz) and high-pass (0.32 Hz) filters were applied. Sleep stages were scored according to Rechtschaffen and Kales (Rechtschaffen and Kales, 1968) using C3 and C4 by two experienced investigators who were blind to the assigned treatment. Differences in scoring between the scorers were resolved by consulting a third experienced investigator.

Off-line detection of slow oscillations was based on an algorithm described in detail previously (Mölle et al., 2002) and was applied to SWS epochs across the entire night. In brief, the EEG for each channel was first low-pass-filtered at 30 Hz and down-sampled to 100 Hz. For the identification of large slow oscillations, a low-pass filter of 3.5 Hz was applied. Then, negative and positive peak potentials were derived from all intervals between consecutive positive-to-negative zero crossings (i.e., one negative and one positive peak for every interval). Only intervals with durations of 0.833–2 seconds (corresponding to a frequency of 0.5–1.2 Hz) were included. We determined each participant’s individual detection thresholds by first calculating the mean values of the negative peak potential and the negative-to-positive peak amplitude of the valid intervals within each channel. These were then averaged across all nine EEG channels. Subsequently, we marked for each channel those intervals as slow oscillation epochs in which the negative peak amplitude was lower than the mean negative peak threshold multiplied by 1.25, and where the amplitude difference (positive peak minus negative peak) was larger than the peak-to-peak amplitude threshold multiplied by 1.25. Negative half-wave peaks were used to characterize slow-oscillation events. Time-frequency plots were derived using the Fieldtrip toolbox for Matlab (http://www.fieldtriptoolbox.org, ‘mtmconvol’ function) on ± 1.5-second windows time-locked to the negative peak of offline-detected slow-oscillation events for frequencies from 5-20 Hz in steps of 0.5 Hz, using a Hanning window corresponding to seven cycles for each frequency, which was further shifted along the time axis in steps of 40 milliseconds. The resulting time-frequency representations were normalized to a pre-event baseline from −1.5 to −1.4 seconds. Statistical comparisons between the mefloquine and placebo conditions were based on two-tailed paired-samples t-tests, which, for correction of multiple comparisons, were applied following a Monte-Carlo-based permutation procedure with 1000 repetitions (resulting alpha-level = 0.05).

#### Serum mefloquine concentrations

Blood was sampled three times (in the evening, in the morning and after the retrieval phase) for the determination of mefloquine concentrations. Blood was centrifuged after 10 minutes of incubation at room temperature and serum samples were frozen at −80°C. Mefloquine levels were measured by high-performance liquid chromatographic analysis with a detection limit of 20ng/ml, an inter-assay coefficient of variation (CV) of 3.28% and an intra-assay CV of 6.52% (Dr. Eberhard & Partner, Dortmund, Germany). Any data points below the detection limit were considered to be 0. Blood samples collected in one placebo session of Experiment 3 were lost but respective values set to 0 because it was this participants’ first experimental session. The third blood sample of the mefloquine session of one individual of Experiment 2 was likewise lost so that this participant’s data were omitted from the analysis of this time point.

#### Control measures: Number encoding, sleepiness, vigilance, mood and general retrieval

During retrieval participants also encoded numbers (Feld et al., 2013) to assess whether mefloquine affects learning performance. Sixteen three-digit numbers were presented on a computer screen for two seconds (500 ms ISI) in three random blocks. After a short break of about one minute, participants were asked to freely recall the numbers and write them on a piece of paper and we analyzed the amount of correct numbers. Afterwards, they were again shown the previously learned numbers, but intermixed with sixteen new numbers, and had to decide which numbers they had already learned. From the hit rate and the false alarm rate we calculated the sensitivity index d-prime to measure recognition memory.

Subjective sleepiness was measured using the one-item Stanford Sleepiness Scale (SSS, Hoddes et al., 1973) during the learning and the retrieval phase as well as in the evening and the morning. Here, participants indicated how sleepy they felt on a scale ranging from one (“Feeling active, vital, alert, or wide awake”) to eight (“Asleep” – provided as an anchor). Objective vigilance was assessed by means of the psycho-motor vigilance task (PVT, Dinges et al., 1997) at the end of the learning and the retrieval phases. In this task, the participants saw a red millisecond counter at the center of a black screen that occasionally and suddenly started counting upwards from 0. Pressing the response button stopped the counter and yielded the reaction time. Participants were asked to react as fast as possible during five minutes and average reaction speed (one divided by reaction time) was used as the objective measure of vigilance (see Table 3 for descriptive data). We assessed the mood of our participants by means of a multidimensional mood questionnaire at four time points per session (Hinz et al., 2012). This questionnaire comprises the three dimensions mood (high values reflect positive mood), tiredness (low values equal tiredness) and calmness (high values reflect calmness). In order to assess whether mefloquine affected general retrieval performance of remote memories that can be expected to be completely consolidated, participants performed a word generation task during retrieval (Regensburg verbal fluency test, Aschenbrenner et al., 2000). Participants were given two minutes each to write down as many words as possible that started with a cue letter (p or m) or belonged to a cue category (hobby or profession). Sum scores of both versions of the task were regarded as a measure of general retrieval performance.

**Table 1.** Mean (standard error of the mean – SEM) performance on the memory tasks, i.e., correctly recalled word-pairs, correctly tapped sequences, accuracy of tapped sequences, correctly recalled card-pair locations and numbers recalled and recognized. Percent of learning refers to absolute score during the Retrieval phase divided by the absolute score during the Learning phase.

|  | Experiment 1 (Sleep) |  |  |  | Experiment 2 (Wake) |  |  |  | Experiment 3 (Retrieval) |  |  |  |
| --- | --- | --- | --- | --- | --- | --- | --- | --- | --- | --- | --- | --- |
|  | Mefloquine |  | Placebo |  | Mefloquine |  | Placebo |  | Placebo |  | Mefloquine |  |
| Word-pairs |  |  |  |  |  |  |  |  |  |  |  |  |
| Learning | 29.11 | (0.93) | 27.83 | (0.80) | 28.75 | (1.08) | 29.17 | (1.01) | 29.83 | (0.89) | 30.00 | (1.10) |
| Retrieval | 28.28 | (1.17) | 29.67 | (1.09) | 27.58 | (1.67) | 27.58 | (1.65) | 29.75 | (1.21) | 30.00 | (1.18) |
| Difference | -0.83 | (0.79) | 1.83 | (0.68) | -1.17 | (1.01) | -1.58 | (1.53) | -0.08 | (1.31) | 0.00 | (0.74) |
| Percent of learning | 0.97 | (0.03) | 1.07 | (0.02) | 0.95 | (0.04) | 0.95 | (0.05) | 1.00 | (0.05) | 1.00 | (0.03) |
| Trials to criterion | 1.50 | (0.19) | 1.67 | (0.16) | 1.75 | (0.18) | 1.67 | (0.14) | 1.33 | (0.19) | 1.33 | (0.14) |
| Finger sequence |  |  |  |  |  |  |  |  |  |  |  |  |
| Correct sequences |  |  |  |  |  |  |  |  |  |  |  |  |
| Learning | 20.79 | (0.91) | 22.18 | (1.02) | 18.64 | (1.33) | 20.67 | (1.32) | 20.58 | (1.55) | 21.75 | (1.58) |
| Retrieval | 25.40 | (1.25) | 25.05 | (1.28) | 23.88 | (1.72) | 24.27 | (1.60) | 25.17 | (2.21) | 25.14 | (2.07) |
| Difference | 4.61 | (0.67) | 2.88 | (0.78) | 5.24 | (0.83) | 3.61 | (0.77) | 4.58 | (0.91) | 3.39 | (1.07) |
| Percent of learning | 1.22 | (0.03) | 1.13 | (0.03) | 1.29 | (0.04) | 1.18 | (0.04) | 1.22 | (0.04) | 1.15 | (0.05) |
| Control sequence | 16.63 | (1.03) | 16.05 | (0.89) | 16.30 | (1.13) | 15.70 | (1.21) | 16.89 | (1.81) | 18.44 | (1.86) |
| Error rates |  |  |  |  |  |  |  |  |  |  |  |  |
| Learning | 0.10 | (0.01) | 0.07 | (0.01) | 0.13 | (0.03) | 0.10 | (0.01) | 0.14 | (0.02) | 0.09 | (0.01) |
| Retrieval | 0.07 | (0.02) | 0.08 | (0.01) | 0.03 | (0.01) | 0.06 | (0.01) | 0.09 | (0.02) | 0.10 | (0.02) |
| Difference | -0.03 | (0.01) | 0.00 | (0.02) | -0.10 | (0.02) | -0.04 | (0.01) | -0.05 | (0.02) | 0.01 | (0.02) |
| Control sequence | 0.14 | (0.02) | 0.14 | (0.03) | 0.05 | (0.01) | 0.12 | (0.02) | 0.14 | (0.04) | 0.11 | (0.02) |
| Numbers |  |  |  |  |  |  |  |  |  |  |  |  |
| Free recall |  |  |  |  |  |  |  |  |  |  |  |  |
| Correct | 7.37 | (0.86) | 8.16 | (0.69) | 7.92 | (1.03) | 6.42 | (0.78) | 7.67 | (0.57) | 6.92 | (0.53) |
| False | 3.95 | (0.86) | 4.05 | (0.57) | 2.83 | (0.61) | 3.75 | (1.08) | 3.00 | (0.49) | 3.17 | (0.49) |
| Percent correct | 0.65 | (0.06) | 0.67 | (0.04) | 0.72 | (0.06) | 0.66 | (0.07) | 0.72 | (0.04) | 0.69 | (0.04) |
| Recognition |  |  |  |  |  |  |  |  |  |  |  |  |
| Hits | 0.78 | (0.04) | 0.85 | (0.03) | 0.76 | (0.05) | 0.67 | (0.05) | 0.72 | (0.04) | 0.69 | (0.04) |
| False alarms | 0.28 | (0.03) | 0.24 | (0.03) | 0.26 | (0.02) | 0.30 | (0.05) | 0.16 | (0.03) | 0.21 | (0.04) |
| d-prime | 1.49 | (0.22) | 1.82 | (0.19) | 1.47 | (0.22) | 1.07 | (0.19) | 1.70 | (0.20) | 1.40 | (0.19) |

**Table 2.** Mean (SEM) of the sleep stages scored according to standard criteria.

|  | **Experiment 1 (Sleep)** |  |  |  | **Experiment 3 (Retrieval)** |  |  |  |
| --- | --- | --- | --- | --- | --- | --- | --- | --- |
|  | Mefloquine |  | Placebo |  | Mefloquine |  | Placebo |  |
| **Percent** |  |  |  |  |  |  |  |  |
| Wake | 3.15 | (0.51) | 3.88 | (0.96) | 3.91 | (1.04) | 1.98 | (0.50) |
| NonREM 1 | 8.03 | (0.59) | 7.37 | (0.53) | 6.51 | (0.89) | 5.57 | (0.60) |
| NonREM 2 | 53.47 | (1.49) | 54.90 | (1.31) | 55.17 | (1.40) | 55.30 | (0.73) |
| NonREM 3 | 9.64 | (0.78) | 8.93 | (0.94) | 8.69 | (0.82) | 9.99 | (0.64) |
| NonREM 4 | 2.53 | (1.24) | 2.19 | (1.04) | 2.64 | (0.67) | 2.29 | (0.72) |
| SWS | 12.18 | (1.40) | 11.12 | (1.41) | 11.33 | (1.22) | 12.29 | (0.88) |
| REM | 22.56 | (1.12) | 22.03 | (1.09) | 21.90 | (1.11) | 23.56 | (1.32) |
| **Minutes** |  |  |  |  |  |  |  |  |
| TST | 463.81 | (4.42) | 461.00 | (5.76) | 471.63 | (2.79) | 471.83 | (2.68) |
| Wake | 14.58 | (2.36) | 17.97 | (4.56) | 18.50 | (4.97) | 9.25 | (2.37) |
| NonREM 1 | 37.06 | (2.65) | 33.78 | (2.35) | 30.71 | (4.18) | 26.33 | (2.88) |
| NonREM 2 | 247.92 | (6.91) | 252.61 | (5.63) | 260.17 | (6.83) | 261.00 | (3.96) |
| NonREM 3 | 44.94 | (3.71) | 41.44 | (4.44) | 41.00 | (3.83) | 47.17 | (3.13) |
| NonREM 4 | 11.97 | (5.90) | 10.33 | (4.93) | 12.50 | (3.14) | 11.00 | (3.46) |
| SWS | 56.92 | (6.75) | 51.78 | (6.68) | 53.50 | (5.71) | 58.17 | (4.36) |
| REM | 104.50 | (5.20) | 101.58 | (5.30) | 103.08 | (5.01) | 110.92 | (5.80) |
| Movement time | 2.83 | (0.35) | 3.28 | (0.34) | 5.67 | (0.72) | 6.17 | (0.52) |

**Table 3:** Mean (SEM) serum mefloquine concentrations (ng/ml), sleepiness (Stanford Sleepiness Scale), reaction speed (psychomotor vigilance task), mood (multi-dimensional mood questionaire) and word generation (verbal fluency test).

|  | Experiment 1 (Sleep) |  |  |  | Experiment 2 (Wake) |  |  |  | Experiment 3 (Retrieval) |  |  |  |
| --- | --- | --- | --- | --- | --- | --- | --- | --- | --- | --- | --- | --- |
|  | Mefloquine |  | Placebo |  | Mefloquine |  | Placebo |  | Mefloquine |  | Placebo |  |
| **Mefloquine levels** |  |  |  |  |  |  |  |  |  |  |  |  |
| Evening | 188.95 | (18.59) | 13.68 | (5.31) | 285.00 | (38.95) | 16.08 | (9.28) | 0.00 | NA | 13.17 | (8.56) |
| Morning | 201.84 | (10.81) | 12.89 | (5.02) | 332.50 | (30.17) | 10.42 | (7.08) | 0.00 | NA | 12.00 | (7.73) |
| Retrieval phase | 207.11 | (12.90) | 13.16 | (5.18) | 312.27 | (29.22) | 11.67 | (8.08) | 354.17 | (32.28) | 11.92 | (7.40) |
| **Sleepiness** |  |  |  |  |  |  |  |  |  |  |  |  |
| Learning phase | 2.58 | (0.25) | 2.68 | (0.28) | 3.00 | (0.43) | 2.50 | (0.26) | 2.46 | (0.24) | 2.83 | (0.27) |
| Evening | 4.26 | (0.29) | 4.32 | (0.32) | 3.58 | (0.45) | 3.25 | (0.39) | 4.21 | (0.29) | 4.50 | (0.38) |
| Morning | 2.68 | (0.25) | 2.84 | (0.24) | 5.42 | (0.43) | 5.42 | (0.36) | 3.00 | (0.3) | 3.33 | (0.26) |
| Retrieval phase | 1.84 | (0.23) | 2.16 | (0.22) | 4.67 | (0.40) | 5.42 | (0.36) | 1.67 | (0.14) | 1.67 | (0.14) |
| **Reaction speed** |  |  |  |  |  |  |  |  |  |  |  |  |
| Learning phase | 3.68 | (0.08) | 3.60 | (0.11) | 3.68 | (0.08) | 3.74 | (0.09) | 3.53 | (0.13) | 3.63 | (0.11) |
| Retrieval phase | 3.78 | (0.09) | 3.69 | (0.10) | 3.58 | (0.13) | 3.40 | (0.15) | 3.71 | (0.16) | 3.63 | (0.13) |
| **Mood** |  |  |  |  |  |  |  |  |  |  |  |  |
| *Positive mood* |  |  |  |  |  |  |  |  |  |  |  |  |
| Learning phase | 4.45 | (0.07) | 4.33 | (0.06) | 4.19 | (0.16) | 4.50 | (0.11) | 4.21 | (0.18) | 4.44 | (0.08) |
| Evening | 4.47 | (0.08) | 4.38 | (0.12) | 4.02 | (0.24) | 4.21 | (0.11) | 4.31 | (0.10) | 4.35 | (0.10) |
| Morning | 4.33 | (0.14) | 4.45 | (0.10) | 3.27 | (0.2) | 3.71 | (0.20) | 3.92 | (0.29) | 4.23 | (0.08) |
| Retrieval phase | 4.54 | (0.15) | 4.49 | (0.12) | 3.85 | (0.24) | 3.48 | (0.31) | 4.65 | (0.06) | 4.58 | (0.09) |
| *Tiredness (inv.)* |  |  |  |  |  |  |  |  |  |  |  |  |
| Learning phase | 3.62 | (0.19) | 3.45 | (0.22) | 3.38 | (0.30) | 3.96 | (0.18) | 3.48 | (0.19) | 3.65 | (0.15) |
| Evening | 2.80 | (0.19) | 2.59 | (0.21) | 3.15 | (0.30) | 3.42 | (0.24) | 2.94 | (0.17) | 2.90 | (0.19) |
| Morning | 3.57 | (0.19) | 3.59 | (0.17) | 2.08 | (0.24) | 2.02 | (0.16) | 3.33 | (0.19) | 3.31 | (0.11) |
| Retrieval phase | 3.97 | (0.17) | 3.67 | (0.21) | 2.40 | (0.27) | 1.94 | (0.23) | 4.19 | (0.15) | 4.17 | (0.16) |
| *Calmness* |  |  |  |  |  |  |  |  |  |  |  |  |
| Learning phase | 4.21 | (0.13) | 4.13 | (0.14) | 3.94 | (0.13) | 4.40 | (0.13) | 3.94 | (0.29) | 4.08 | (0.17) |
| Evening | 4.22 | (0.11) | 4.03 | (0.18) | 3.90 | (0.2) | 4.25 | (0.16) | 4.25 | (0.15) | 4.38 | (0.09) |
| Morning | 4.21 | (0.13) | 4.08 | (0.11) | 3.44 | (0.17) | 3.79 | (0.15) | 3.75 | (0.23) | 4.27 | (0.10) |
| Retrieval phase | 4.11 | (0.17) | 4.04 | (0.21) | 3.63 | (0.18) | 3.54 | (0.23) | 4.31 | (0.15) | 4.33 | (0.17) |
| **Word generation** |  |  |  |  |  |  |  |  |  |  |  |  |
| Letter | 16.16 | (1.05) | 17.37 | (1.16) | 14.92 | (1.41) | 13.92 | (1.02) | 14.58 | (1.58) | 14.25 | (1.30) |
| Category | 20.42 | (0.84) | 21.84 | (0.98) | 17.83 | (1.30) | 15.58 | (1.25) | 19.58 | (0.95) | 17.67 | (1.30) |
| Total | 36.58 | (1.64) | 39.21 | (1.80) | 32.75 | (2.45) | 29.50 | (1.86) | 34.17 | (2.10) | 31.92 | (2.17) |

#### Statistical analysis

One participant of Experiment 1 was excluded from data analyses because of insufficient sleep in the placebo night. Word-pair data of another participant of Experiment 1 were discarded because of retention performance of more than 2 standard deviations below the group mean. Finger sequence tapping data of one participant of Experiment 2 were excluded because of extreme underperformance during the last three blocks of the learning phase (compared to the rest of the learning) in one condition reflecting underrepresentation of encoding. Psychomotor vigilance task data of two participants (from Experiments 2 and 3) were discarded because they indicated a misconception of the instructions (repeated responses in the absence of cues); one time point of assessment was lost in Experiment 1 due to technical malfunction.

Statistical analyses generally relied on analyses of covariance. Since mefloquine was detectable in samples collected in the placebo session of six, three and three participants in Experiments 1, 2 and 3, respectively, which constitutes roughly 50% of cases in which the placebo session took place after the mefloquine session (see Table 3 for descriptive data), serum mefloquine values during the learning phase of the placebo session were included as covariates. (The main conclusion of the study, i.e., reduced declarative memory consolidation under mefloquine, remains if the statistical analyses are conducted without the covariate.) Greenhouse-Geisser correction of degrees of freedom was performed whenever necessary. Control measures were compared by paired t-tests at each time point to detect intra-experimental differences unless specified otherwise.

## Results

### Mefloquine concentrations

The timing of mefloquine administrations in the three experiments worked as planned. Serum mefloquine concentrations were strongly increased in the mefloquine as compared to the placebo condition in Experiments 1 (sleep) and 2 (wake) at all time points (all p ≤ 0.001), as well as in Experiment 3 at retrieval (t_(11)_ = 10.21, p ≤ 0.001, all other time points p ≥ 0.15).

### Declarative word-pair memory

Regarding declarative memory, there were no differences in any of the experiments in encoding performance measured as the number of word-pairs correctly recalled during learning (all p ≥ 0.18, see Table 1 for descriptive data). Mefloquine compared to placebo impaired retention of the declarative word-pair task in sleeping subjects only (mefloquine: −0.83 (0.79), placebo: 1.83 (0.68) word-pairs, F_(1,16)_ = 29.75, p ≤ 0.001, η_p_^2^ = 0.65; Figure 1B). In contrast, this effect was not observed in the wake control (mefloquine: − 1.17 (1.00), placebo: −1.58 (1.53), F(1,10) = 2.43, p = 0.15, η_p_^2^ = 0.20) and retrieval control groups (mefloquine: −0.08 (1.31), placebo: 0.00 (0.74), F(1,10) = 0.14, p = 0.72, η_p_^2^ = 0.014). This pattern was statistically corroborated by an overall interaction effect between the experimental groups regarding treatment (F(1,39) = 4.52, p = 0.040, η_p_^2^ = 0.10), meaning that mefloquine reduced retention performance for word-pairs exclusively in the sleep experiment. The beneficial effect on declarative memory retention of sleep per se was confirmed in a comparison covering only the placebo conditions of the experiments (F_(1,39)_ = 4.41, p = 0.042, η_p_^2^ = 0.10), which also indicates that the sleep effect was not affected by the long learning-to-sleep delay used here.

### Procedural fingers sequence memory

In the procedural memory domain, encoding performance of finger sequence tapping, i.e., the amount of correctly tapped sequences in the last three blocks of the learning phase, was comparable across experiments and conditions (all p ≥ 0.14, see Table 1 for descriptive data). Overall analyses revealed a mefloquine-induced improvement of finger sequence retention, i.e., more correctly tapped sequences, that emerged irrespective of the experimental group (main effect of treatment, F_(1,38)_ = 3.97, p = 0.054, η_p_^2^ = 0.10; treatment × experiment, F_(2,38)_ = 0.09, p = 0.91, η_p_^2^ = 0.01). Performance accuracy as reflected by error rates, i.e., incorrectly tapped sequences divided by total tapped sequences, showed an even more pronounced general improvement across the retention interval under mefloquine (main effect of treatment, F_(1,38)_ = 8.74, p = 0.005, η_p_^2^ = 0.19; treatment × experiment, F_(2,38)_ = 0.65, p = 0.53, η_p_^2^ = 0.03). Comparing retention performance between the placebo conditions, we did not observe a beneficial effect of sleep on tapping performance (F_(1,39)_ = 0.22, p = 0.65, η_p_^2^ = 0.01). We found some evidence that mefloquine enhanced reaction times in a control vigilance task (wake experiment: t_(10)_ = 2.30, p = 0.044; see *Control measures* for details), possibly supporting procedural execution independent of memory processes. However, this explanation is not supported by the fact that performance on a control finger sequence, tested exclusively at retrieval, remained unaffected by the drug (all p ≥ 0.12 for number of correct sequences and error rate).

### Sleep architecture and sleep oscillations

Mefloquine administration before sleep in Experiment 1 had no effect on gross sleep architecture (all p ≥ 0.23, see Figure 2A, and Table 2 for descriptive data). Likewise, no differences between the sleep conditions were observed, when mefloquine was administered after sleep in Experiment 3 (all p ≥ 0.16). In more fine-grained analyses of the results of Experiment 1 we explored potential mefloquine effects on the relationship between slow oscillations (0.75 Hz) and sleep spindles, i.e., the most prominent features of NonREM sleep to benefit memory function (Clemens et al., 2007; Staresina et al., 2015). Mefloquine reduced power in the fast spindle band (12-15 Hz) during the up-state of the slow oscillation and, moreover, decreased power in the slow spindle band (9-12 Hz) during the respective up- to down-state transition (see Figure 2B for descriptive and statistical data).

**Figure 2.**
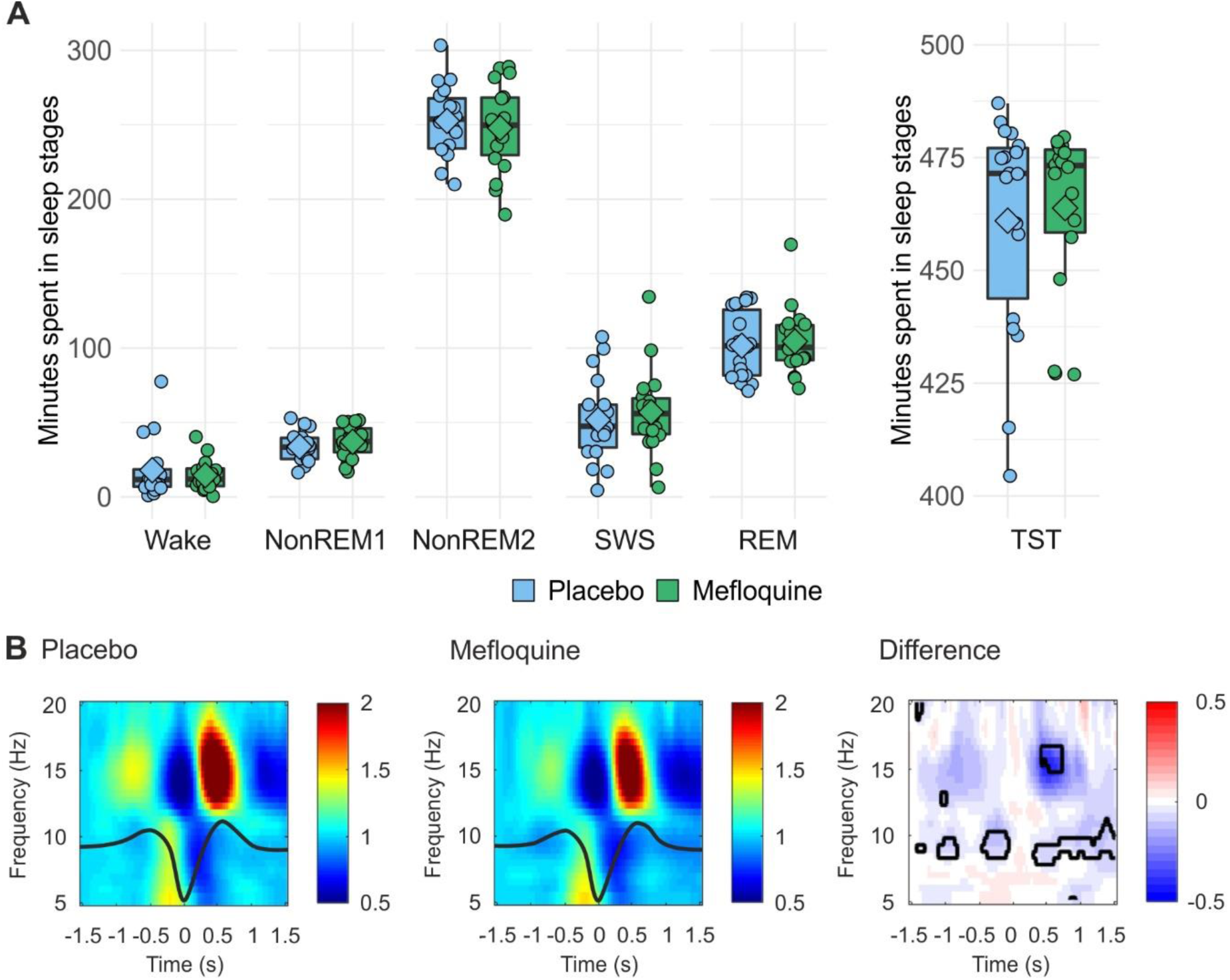
Sleep results of Experiment 1. (A) Box and whisker plot (thick line: median, hinge: 25% and 75% quartiles, whiskers: smallest/largest values no further than 1.5 * inter quartile range from the hinge), mean (diamond shape) and individual data points (circles) of time spent in different sleep stages (SWS – slow wave sleep, REM – rapid eye movement sleep, TST – total sleep time) for the mefloquine condition (green) and the placebo condition (blue) scored according to standard criteria (for details see methods and for sleep results of Experiment 3 see SI results). Mefloquine did not induce statistically significant differences in overall sleep architecture, indicating that the drug-induced declarative memory impairment observed in this experiment cannot be attributed to gross sleep disturbances. (B) Effects of mefloquine on the coupling of sleep spindles to slow oscillations (0.75 Hz) were analyzed by identifying the peaks of the slow oscillation down-states and calculating EEG power between 5 and 20 Hz in 3-second time windows centered on the down-states (x-axis: ‘0’). The left and middle panels depict this time-frequency representation for the placebo and the mefloquine conditions with the grand average of the detected slow oscillations overlaid as thick solid lines (slow oscillations are derived from EEG channel Pz; see Extended Data Figure 1-1 for Fz and Cz). The right panel depicts the difference between the two conditions, with the areas of statistically significant differences outlined in black (two-tailed paired-samples t-tests corrected for multiple comparisons). Power in the upper fast spindle band around 15 Hz was reduced by mefloquine during the up-state of the slow oscillation, and power in the slow spindle band (9-12 Hz) was reduced by the drug in the up- to down-state transition. Thus, mefloquine suppressed oscillatory coupling during sleep, an effect that may have mediated the drug-induced impairment of declarative memory retention.

### Control measures: Number encoding, sleepiness, vigilance, mood and general retrieval

For number encoding we used mefloquine levels during the retrieval (rather than encoding) phase as covariates. Comparing sleep to wakefulness using only data from the placebo conditions, we found an enhancing effect of sleep on recognition (F_(1,40)_ = 6.06, p = 0.018, η_p_^2^ = 0.13) but not on free recall (F_(1,40)_ = 1.88, p = 0.178, η_p_^2^ = 0.05, see Table 1 for descriptive data). Also, there was an improvement in recognition under mefloquine in the wake experiment (F_(1,10)_ = 7.20, p = 0.023, η_p_^2^ = 0.42) that was not evident in the sleep (F_(1,17)_ = 0.01, p = 0.91, η_p_^2^ ≤ 0.01) or retrieval experiment (F(1,10) = 1.79, p = 0.210, ηp2 = 0.15), which was reflected in a trend-wise respective interaction (main effect of treatment: F_(1,39)_ = 4.56, p = 0.039, η_p_^2^ = 0.11, treatment × experiment: F_(2,39)_ = 3.02, p = 0.060, η_p_^2^ = 0.13).

In Experiment 2 (nocturnal wakefulness), a trend towards less subjective sleepiness at retrieval emerged in the mefloquine condition (t_(11)_ = −1.83, p = 0.095, all other p ≥ 0.11). As expected, when comparing the experiments by ANOVA (factors: treatment, sleep/wake and time points) we found that staying awake increased subjective sleepiness (sleep/wake × time point interaction: F_(3,123)_ = 46.41, p ≤ 0.001, η_p_^2^ = 0.53). Follow-up comparisons verified that this effect focused on the morning and retrieval phase assessments in both treatment conditions (all p ≤ 0.001).

In accordance with subjective sleepiness, in Experiment 2, there was a significant mefloquine-induced improvement in reaction speed during retrieval as measured by the psychomotor vigilance task (t_(10)_ = 2.30, p = 0.044, all other p ≥ 0.13). Here, comparing the experiments revealed a treatment × time-point interaction (F_(3,38)_ = 8.79, p = 0.005, η_p_^2^ = 0.19) that was mainly driven by increased reaction speed under mefloquine at retrieval. As expected, ANOVA indicated that reaction speed was generally reduced during retrieval in the wake compared to the sleep groups (sleep/wake × time point: F_(3,38)_ = 25.85, p ≤ 0.001, η_p_^2^ = 0.41).

An ANOVA indicated that mood was negatively affected by sleep deprivation (sleep/wake × time point: F_(3,123)_ = 10.30, p ≤ 0.001, η_p_^2^ = 0.20), as well as a time-dependent influence of mefloquine on mood (treatment × time point: F_(3,123)_ = 6.29, p ≤ 0.001, η_p_^2^ = 0.13), however, this interaction could not be pin-pointed to any specific direction and/or time point. Tiredness was increased by sleep deprivation at the expected time points (sleep/wake × time point: F_(3,123)_ = 45.76, p ≤ 0.001, η_p_^2^ = 0.53) and mefloquine reduced tiredness mainly in the retrieval phase (treatment × time point: F_(3,123)_ = 3.38, p = 0.022, η_p_^2^ = 0.08). Similarly, sleep deprivation reduced calmness at the two later time points (sleep/wake × time point: F_(3,123)_ = 5.65, p = 0.002, η_p_^2^ = 0.12).

Regarding general retrieval performance, there were no differences between mefloquine and placebo in any of the experiments (all p ≥ 0.10). Comparing the individual experiments, however, we found a main impairing effect of sleep deprivation (F_(1,41)_ = 4.78, p = 0.035, η_p_^2^ = 0.10).

## Discussion

In the current study, we administered mefloquine to block gap junctions during nocturnal sleep after participants learned a declarative word-pair task and a procedural finger-sequence tapping task. As hypothesized, blocking gap junctions impaired the beneficial effect of sleep on declarative memory consolidation in the main experiment, whereas administering the drug before nocturnal wakefulness or at retrieval testing did not affect declarative memory. Surprisingly, mefloquine improved retention of the procedural finger sequence tapping task across all three experiments. We did not find an effect of mefloquine on neurophysiological sleep architecture, but detected a marked mefloquine-induced disruption of sleep spindle-to-slow oscillation coupling. We conclude that mefloquine specifically disrupted mechanisms of consolidation that strengthen declarative memory formation during sleep, but did not influence respective processes during wakefulness, nor retrieval function. These results suggest that mefloquine effectively blocked neuro-electric processes of memory replay that rely on gap junctions to produce fast network oscillations (Maier et al., 2003; Diba and Buzsaki, 2007) and are at the core of time-compressed sleep-dependent memory consolidation (Ji and Wilson, 2007; Diekelmann and Born, 2010).

The uncoupling of spindles and slow oscillations by mefloquine points toward an impairing effect of the drug on the interplay between replay and spindle generation. Fast spindle density increases with the number of word-pairs encoded before sleep (Gais et al., 2002), and boosting up-state-associated fast spindle activity by auditory stimulation enhances word-pair retention across sleep (Ngo et al., 2013). It is therefore tempting to speculate that blocking hippocampal replay via mefloquine not only impaired declarative memory retention but also reduced corresponding slow oscillation up-state-associated spindle activity, a notion that is consistent with evidence suggesting a direct impact of hippocampal ripples on spindle activity (Peyrache et al., 2011). However, the direct measurement of sharp-wave/ripples is not feasible in healthy human subjects.

Importantly, the results in the procedural domain strongly indicate that the detrimental effect of mefloquine on sleep-specific consolidation mechanisms is restricted to declarative memory formation that has been most closely linked to replay via sharp-wave/ripples in the hippocampus. Although cued reactivation has also been shown to enhance procedural memory retention during sleep (Antony et al., 2012) this has not been directly linked to hippocampal replay. Therefore, our seemingly paradoxical finding of increased procedural retention can be plausibly explained by a mefloquine-induced reduction in hippocampal-striatal competition, since such a reduction has been shown to facilitate procedural memory consolidation during post-learning sleep (Albouy et al., 2013). The absence of a clear consolidation benefit of sleep for procedural memory may have been due to the longer-than-usual period of wake retention scheduled before sleep (Nettersheim et al., 2015), which we were forced employ to enable optimal uptake of the drug, suggesting that this task is less robust against extended intervals between learning and sleep. The increased reaction speed in the vigilance control task we found at retrieval may indicate that enhanced retention performance on the finger sequence tapping task under mefloquine was mediated by increased reaction speed. However, the fact that we observed no such improvement for the control finger sequence and that the error rate was equally reduced for the retained sequence makes this account unlikely.

Our finding of increased learning of numbers after sleep compared to wakefulness is in line with previous reports of improved encoding after sleep (Mander et al., 2011). However, we could not find the previously reported detrimental effect of mefloquine on encoding (van Essen et al., 2010; Bissiere et al., 2011) and, in the wake group, an increase in encoding performance was even evident. This effect possibly reflects arousing effects of mefloquine, as likewise suggested by control measures (see below), which can be expected to be most prominent after sleep deprivation. As our experiment was not primarily designed to test encoding, its results should not be interpreted to refute previous reports of mefloquine-induced impairments in memory acquisition in rats (Bissiere et al., 2011). The pattern of results of subjectively reduced sleepiness and tiredness as well as objectively enhanced vigilance in the control measures after mefloquine intake especially in the wake group hints at an unspecific arousal effect of the drug that becomes most evident after sleep deprivation.

Our main results of reduced retention of word-pair memory under mefloquin during sleep identify an as yet unknown mechanism of sleep-dependent memory formation. They put a new complexion on the “neuron doctrine,” i.e., the concept of the brain as a composite of discrete individual neurons that communicate exclusively via neurochemicals, which was first and foremost touted by Santiago Ramón y Cajal (Yuste, 2015). This assumption was contested as early as the nineteen-fifties (Bullock, 1959) and it is now clear that the direct electrical coupling of neurons via gap junctions indeed represents a major mode of neuronal information exchange (Bennett and Zukin, 2004; Sohl et al., 2005). Recent studies in rats indicate that gap junctions support acquisition and maintenance of fear memories (Bissiere et al., 2011). In those experiments, which did not manipulate sleep in the retention interval, intraperitoneal and intra-hippocampal administration of gap junction blockers (mefloquine and carbenoxolone) before or directly after learning blocked the subsequent retrieval of context fear memories, while the injection immediately preceding retrieval had no effect. Of note, blocking gap junctions during learning also inhibits hippocampal theta oscillations, which are known to support encoding (Fell and Axmacher, 2011). In humans, blocking gap junctions so far has been shown to impair low-level learning, i.e., the acquisition of eye blink conditioning (van Essen et al., 2010), but evidence in the high-level declarative memory domain was lacking.

A main strength of our experimental approach is the use of human participants, which allows direct interpretation of the results without translating them from an animal model. However, this approach also constrains the amount of control that is feasible. While mefloquine is a potent blocker of connexin 36, the primary gap junction protein in electrical synapses of the mammalian central nervous system, it also has effects that are unrelated to gap junctions (Cruikshank et al., 2004). Nevertheless, its effects on memory consolidation found in rats, i.e., freezing in a fear-conditioned context was reduced, if mefloquine was given after training, where identical to those of the general gap-junction blocker carbonoxolone that has a different spectrum of non-gap junction-related effects (Bissiere et al., 2011). We also cannot rule out that the observed effect may be related to the suppression of theta oscillations within the hippocampus, e.g., during REM sleep, that has been described during awake learning in rats (Bissiere et al., 2011). Future work in animal models will undoubtedly shed more light onto such alternatives.

In sum, we demonstrate in humans that blocking gap junctions during nocturnal sleep selectively impairs declarative word-pair retention, an effect presumably mediated via inhibition of sharp-wave/ripples and the associated time-compressed replay of memory traces (Girardeau et al., 2009; van de Ven et al., 2016). These data indicate a functional role for electrical synapses in the sleep-dependent consolidation of high-level declarative memory in humans. Concurrent sleep and declarative memory dysfunction is a common feature of cognitive impairments that hallmark ailments such as schizophrenia (Manoach et al., 2016) and Alzheimer’s disease (Lim et al., 2013), but also emerge during normal aging (Mander et al., 2013; Helfrich et al., 2018). Given that gap junctions have also been suggested to play a role both in schizophrenia (Aleksic et al., 2007) and Alzheimer’s disease (Giaume et al., 2019), unraveling their role in sleep-dependent memory formation in humans may open new therapeutic avenues beyond the manipulation of chemical neuroreceptors, which all in all has not produced many recent pharmacological innovations (MacEwan et al., 2016; Feld and Born, 2019). Electrical synapses might even be targeted in combination with the external cueing of memory traces during sleep to erase maladaptive behavior and/or strengthen beneficial alternatives (Hauner et al., 2013). Moreover, our result of sleep-associated mnemonic impairments following mefloquine intake may also shed new light on reports that mefloquine regimens for malaria prevention can trigger adverse neuropsychiatric effects (Chen et al., 2007; Saunders et al., 2015).

## Acknowledgments

This work was supported by grants from the Deutsche Forschungsgemeinschaft (DFG; SFB 654 “Plasticity and Sleep”) and from the German Federal Ministry of Education and Research (BMBF) to the German Center for Diabetes Research (DZD e.V.; 01GI0925), as well as an Emmy Noether Fellowship to G.B.F. (FE 1617/2-1).

## Author contributions

G.B.F., M.H., and J.B. designed the study and wrote the manuscript. G.B.F., M.H. and H.V.N. analysed and interpreted the data. S.G., L.K. and K.B. collected and analysed the data under the supervision of G.B.F. A.F. was the medical supervisor and consultant of the study on all pharmacology-related issues. E.D., H.V.N., A.F., S.G., L.K. and K.B. critically advised on the manuscript and interpretation of the data.

## Extended Data

### Polysomnographical frequency analyses

As outlined in the main text, we found a reduction at Pz in slow and fast spindle power during the down- and up-state of the slow-oscillation, respectively. Supplementary Figure S1 details descriptive and inference statistics for the all three central EEG channels Fz, Cz and Pz).

**Supplementary Figure 2-1.**
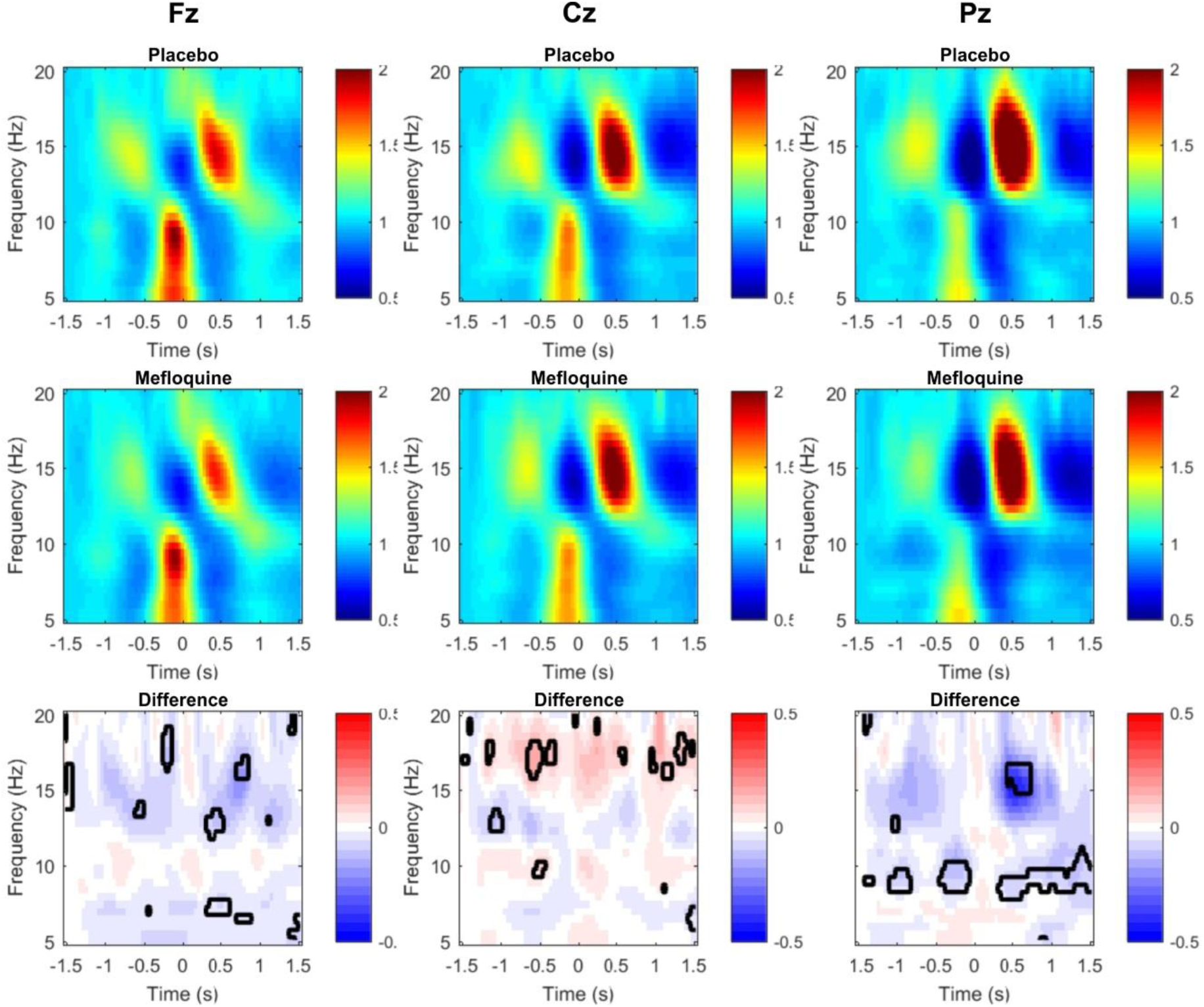
Time-frequency plots time-locked to the down-state of the slow oscillation during SWS. Top panels depict the placebo and midline panels the mefloquine conditions. Bottom panels show the difference between conditions as well as statistical significance based on two-tailed paired-samples t-test with a Monte-Carlo-based permutation procedure with 1000 repetitions and an alpha-level set to 0.05 (this procedure corrects for multiple comparisons).

